# Genetic Architecture of Subcortical Brain Structures in Over 40,000 Individuals Worldwide

**DOI:** 10.1101/173831

**Authors:** Claudia L Satizabal, Hieab HH Adams, Derrek P Hibar, Charles C White, Jason L Stein, Markus Scholz, Murali Sargurupremraj, Neda Jahanshad, Albert V Smith, Joshua C Bis, Xueqiu Jian, Michelle Luciano, Edith Hofer, Alexander Teumer, Sven J van der Lee, Jingyun Yang, Lisa R Yanek, Tom V Lee, Shuo Li, Yanhui Hu, Jia Yu Koh, John D Eicher, Sylvane Desrivières, Alejandro Arias-Vasquez, Ganesh Chauhan, Lavinia Athanasiu, Miguel E Renteria, Sungeun Kim, David Höhn, Nicola J Armstrong, Qiang Chen, Avram J Holmes, Anouk den Braber, Iwona Kloszewska, Micael Andersson, Thomas Espeseth, Oliver Grimm, Lucija Abramovic, Saud Alhusaini, Yuri Milaneschi, Martina Papmeyer, Tomas Axelsson, Stefan Ehrlich, Roberto Roiz-Santiañez, Bernd Kraemer, Asta K Håberg, Hannah J Jones, G Bruce Pike, Dan J Stein, Allison Stevens, Janita Bralten, Meike W Vernooij, Tamara B Harris, Irina Filippi, A Veronica Witte, Tulio Guadalupe, Katharina Wittfeld, Thomas H Mosley, James T Becker, Nhat Trung Doan, Saskia P Hagenaars, Yasaman Saba, Gabriel Cuellar-Partida, Najaf Amin, Saima Hilal, Kwangsik Nho, Nazanin Karbalai, Konstantinos Arfanakis, Diane M Becker, David Ames, Aaron L Goldman, Phil H Lee, Dorret I Boomsma, Simon Lovestone, Sudheer Giddaluru, Stephanie Le Hellard, Manuel Mattheisen, Marc M Bohlken, Dalia Kasperaviciute, Lianne Schmaal, Stephen M Lawrie, Ingrid Agartz, Esther Walton, Diana Tordesillas-Gutierrez, Gareth E Davies, Jean Shin, Jonathan C Ipser, Louis N Vinke, Martine Hoogman, Maria J Knol, Tianye Jia, Ralph Burkhardt, Marieke Klein, Fabrice Crivello, Deborah Janowitz, Owen Carmichael, Unn K Haukvik, Benjamin S Aribisala, Helena Schmidt, Lachlan T Strike, Ching-Yu Cheng, Shannon L Risacher, Benno Pütz, Debra A Fleischman, Amelia A Assareh, Venkata S Mattay, Randy L Buckner, Patrizia Mecocci, Anders M Dale, Sven Cichon, Marco P Boks, Mar Matarin, Brenda WJH Penninx, Vince D Calhoun, M Mallar Chakravarty, Andre Marquand, Christine Macare, Shahrzad Kharabian Masouleh, Jaap Oosterlaan, Philippe Amouyel, Katrin Hegenscheid, Jerome I Rotter, Andrew J Schork, David CM Liewald, Greig I De Zubicaray, Tien Yin Wong, Li Shen, Philipp G Sämann, Henry Brodaty, Joshua L Roffman, Eco JC De Geus, Magda Tsolaki, Susanne Erk, Kristel R Van Eijk, Gianpiero L Cavalleri, Nic JA Van der Wee, Andrew M McIntosh, Randy L Gollub, Kazima B Bulayeva, Manon Bernard, Jennifer S Richards, Jayandra J Himali, Markus Loeffler, Nanda Rommelse, Wolfgang Hoffmann, Lars T Westlye, Maria C Valdés Hernández, Narelle K Hansell, Theo GM Van Erp, Christiane Wolf, John BJ Kwok, Bruno Vellas, Andreas Heinz, Loes M Olde Loohuis, Norman Delanty, Beng-Choon Ho, Christopher RK Ching, Elena Shumskaya, Albert Hofman, Dennis Van der Meer, Georg Homuth, Bruce M Psaty, Mark Bastin, Grant W Montgomery, Tatiana M Foroud, Simone Reppermund, Jouke-Jan Hottenga, Andrew Simmons, Andreas Meyer-Lindenberg, Wiepke Cahn, Christopher D Whelan, Marjolein MJ Van Donkelaar, Qiong Yang, Norbert Hosten, Robert C Green, Anbupalam Thalamuthu, Sebastian Mohnke, Hilleke E Hulshoff Pol, Honghuang Lin, Clifford R Jack, Peter R Schofield, Thomas W Mühleisen, Pauline Maillard, Steven G Potkin, Wei Wen, Evan Fletcher, Arthur W Toga, Oliver Gruber, Matthew Huentelman, George Davey Smith, Lenore J Launer, Lars Nyberg, Erik G Jönsson, Benedicto Crespo-Facorro, Nastassja Koen, Douglas Greve, André G Uitterlinden, Daniel R Weinberger, Vidar M Steen, Iryna O Fedko, Nynke A Groenewold, Wiro J Niessen, Roberto Toro, Christophe Tzourio, William T Longstreth, M Kamran Ikram, Jordan W Smoller, Marie-Jose Van Tol, Jessika E Sussmann, Tomas Paus, Hervé Lemaître, Bernard Mazoyer, Ole A Andreassen, Florian Holsboer, Dick J Veltman, Jessica A Turner, Zdenka Pausova, Gunter Schumann, Daan Van Rooij, Srdjan Djurovic, Ian J Deary, Katie L McMahon, Bertram Müller-Myhsok, Rachel M Brouwer, Hilkka Soininen, Massimo Pandolfo, Thomas H Wassink, Joshua W Cheung, Thomas Wolfers, Jean-Luc Martinot, Marcel P Zwiers, Matthias Nauck, Ingrid Melle, Nicholas G Martin, Ryota Kanai, Eric Westman, René S Kahn, Sanjay M Sisodiya, Tonya White, Arvin Saremi, Hans van Bokhoven, Han G Brunner, Henry Völzke, Margaret J Wright, Dennis Van 't Ent, Markus M Nöthen, Roel A Ophoff, Jan K Buitelaar, Guillén Fernández, Perminder S Sachdev, Marcella Rietschel, Neeltje EM Van Haren, Simon E Fisher, Alexa S Beiser, Clyde Francks, Andrew J Saykin, Karen A Mather, Nina Romanczuk-Seiferth, Catharina A Hartman, Anita L DeStefano, Dirk J Heslenfeld, Michael W Weiner, Henrik Walter, Pieter J Hoekstra, Paul A Nyquist, Barbara Franke, David A Bennett, Hans J Grabe, Andrew D Johnson, Christopher Chen, Cornelia M van Duijn, Oscar L Lopez, Myriam Fornage, Joanna A Wardlaw, Reinhold Schmidt, Charles DeCarli, Philip L De Jager, Arno Villringer, Stéphanie Debette, Vilmundur Gudnason, Sarah E Medland, Joshua M Shulman, Paul M Thompson, Sudha Seshadri, M Arfan Ikram

## Abstract

Subcortical brain structures are integral to motion, consciousness, emotions, and learning. We identified common genetic variation related to the volumes of nucleus accumbens, amygdala, brainstem, caudate nucleus, globus pallidus, putamen, and thalamus, using genome-wide association analyses in over 40,000 individuals from CHARGE, ENIGMA and the UK-Biobank. We show that variability in subcortical volumes is heritable, and identify 25 significantly associated loci (20 novel). Annotation of these loci utilizing gene expression, methylation, and neuropathological data identified 62 candidate genes implicated in neurodevelopment, synaptic signaling, axonal transport, apoptosis, and susceptibility to neurological disorders. This set of genes is significantly enriched for *Drosophila* orthologs associated with neurodevelopmental phenotypes, suggesting evolutionarily conserved mechanisms. Our findings uncover novel biology and potential drug targets underlying brain development and disease.

---

Subcortical brain structures are essential for the control of autonomic and sensorimotor functions^1,2^, modulation of processes involved in learning, memory, and decision-making^3,4^, as well as in emotional reactivity^5,6^ and consciousness^7^. They often act through networks influencing input to and output from the cerebral cortex^8,9^. The pathology of many cognitive, psychiatric, and movement disorders is restricted to, begins in, or predominantly involves subcortical brain structures and related circuitries^10^. For instance, tau pathology has shown to manifest itself early in the brainstem and thalamic nuclei of individuals with Alzheimer’s disease before spreading to cortical areas through efferent networks^11^. Similarly, the formation of Lewy bodies and Lewy neurites in Parkinson’s disease appears early in the lower brainstem (and olfactory structures) before affecting the substantia nigra^12^.

A recent investigation identified five novel genetic loci influencing the volumes of the putamen and caudate, which pointed to genes controlling neuronal growth, apoptosis, and learning^13^. However, no genome-wide significant signals associated with the volumes of the nucleus accumbens, amygdala, globus pallidus, and thalamus were detected, and the genetic variation associated with brainstem volume has not been previously explored. Identifying novel genetic factors contributing to variability in subcortical structures, including the brainstem, should further improve our understanding of brain development and disease.

We sought to identify novel genetic variants influencing the volumes of seven subcortical structures (nucleus accumbens, amygdala, caudate nucleus, putamen, globus pallidus, thalamus, and brainstem (including mesencephalon, pons, and medulla oblongata)), through genome-wide association (GWA) analyses in over 40,000 individuals from 54 study samples (Table S1) from the Cohorts of Heart and Aging Research in Genomic Epidemiology (CHARGE) consortium, the Enhancing Neuro Imaging Genetics through Meta-Analysis (ENIGMA) consortium, and the United Kingdom Biobank (UKBB).

## RESULTS

### Heritability

To examine the extent to which genetic variation accounts for variation in subcortical brain volumes, we estimated the heritability of those volumes in the Framingham Heart Study (FHS) and the Austrian Stroke Prevention Study (ASPS-Fam) family-based cohorts. Our analyses are in line with previous studies conducted in young^14^ and older^15^ twins, suggesting that variability in subcortical volumes is moderately to highly heritable. The structures with highest heritability in the FHS and the ASPS-Fam family-based cohorts are the brainstem (ranging from 79-86%), caudate nucleus (71-85%), putamen (71-79%) and nucleus accumbens (66%); followed by the globus pallidus (55-60%), thalamus (47-54%), and amygdala (34-59%) (Figure 1 and Supplementary Table S2).

**Figure 1.**
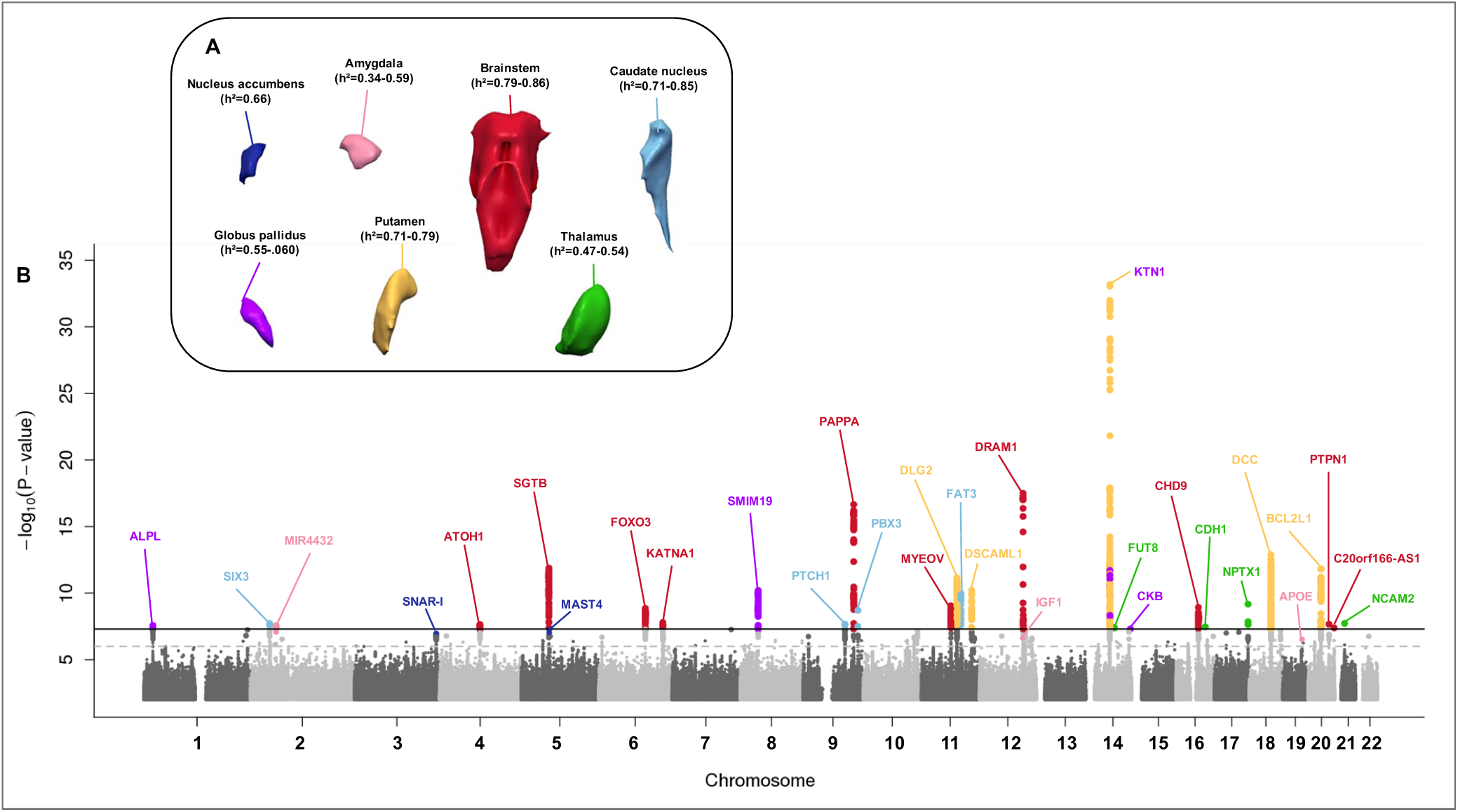
Heritability and Manhattan plot of genetic variation associated with subcortical brain volumes in the discovery sample. Analyses were adjusted for sex, age, age^2^, total intracranial volume, and population structure. **A.** Heritability (h^2^) estimates were performed with SOLAR in the Framingham Heart Study (n=895) and the Austrian Stroke Prevention-Family Study (n=370). **B.** Combined Manhattan plot. Each dot denotes a single genetic variant plotted according to its genomic position (x-axis) and -loglO(P) for the associations with each subcortical volume (y-axis). Variants are colored differently for each structure (see legend in A). The solid horizontal line denotes genome-wide significance (P < 5 x 10 ^-8^), the dashed horizontal line denotes a threshold of P < 10^-6^. Individual Manhattan plots may be found in the Supplementary note.

### Genome-wide associations

We undertook a GWA analysis on the MRI-derived volumes of subcortical structures using the 1000 Genomes Project^1516^ reference panel (phase 1 v.3) for imputation of missing variants. Our discovery sample comprised up to n=25,587 individuals of European ancestry from 45 study samples in CHARGE and ENIGMA (Table S1). Additionally, we included four samples for replication in Europeans (up to n=13,707), three for generalization to African-Americans (up to n=769), and two for generalization to Asians (n=341). Each study related genetic variants with minor allele frequency (MAF) ≥1% to the volumes of subcortical structures (average volume for bilateral structures) using additive genetic models adjusted for sex, age, age^2^, total intracranial volume (total brain volume in the UKBB), and population structure. After quality control, we combined study-specific GWA results using sample-size-weighted fixed effects methods in METAL^16^. We conducted meta-analyses in stages, from discovery, through replication and generalization, to the combination of all available samples.

In the discovery analysis, we identified 25 genome-wide significant loci across six subcortical structures, 20 of which are novel (Table 1). Among them, 13 variants were located within genes (one 3’-UTR, one missense, one non-coding transcript, 10 intronic), and 12 in intergenic regions. In addition to these 25 loci, a further seven novel probable genetic associations were identified: four had p-values just above the threshold of significance (5.3 x 10^-8^ to 2.9 x 10 ^-7^) and three others reached genome-wide significance but were less frequent variants reliably genotyped in a smaller sample of n <2500 individuals. Replication results in the UKBB are shown in Table 1. We carried forward these 32 loci pointing to 31 candidate genes (variants at the 14q22.3 locus near KTN1 were related to putamen and globus pallidus volumes) to *in-silico* replication in Europeans, generalization in African-Americans and Asians, and combined meta-analysis of all samples (Table S3). Of 32 candidate loci, the direction of association was the same for 24 variants in Europeans and 15 variants across all ethnicities. In the combined meta-analysis, 21 of the 32 associations were genome-wide significant, 20 for which the strength of association increased from the discovery. Among these, are 2 of variants for the nucleus accumbens (*MAST4* and *SNAR-I)* below the threshold in the discovery now reached genome-wide significance in the combined meta-analysis.

**Table 1.**
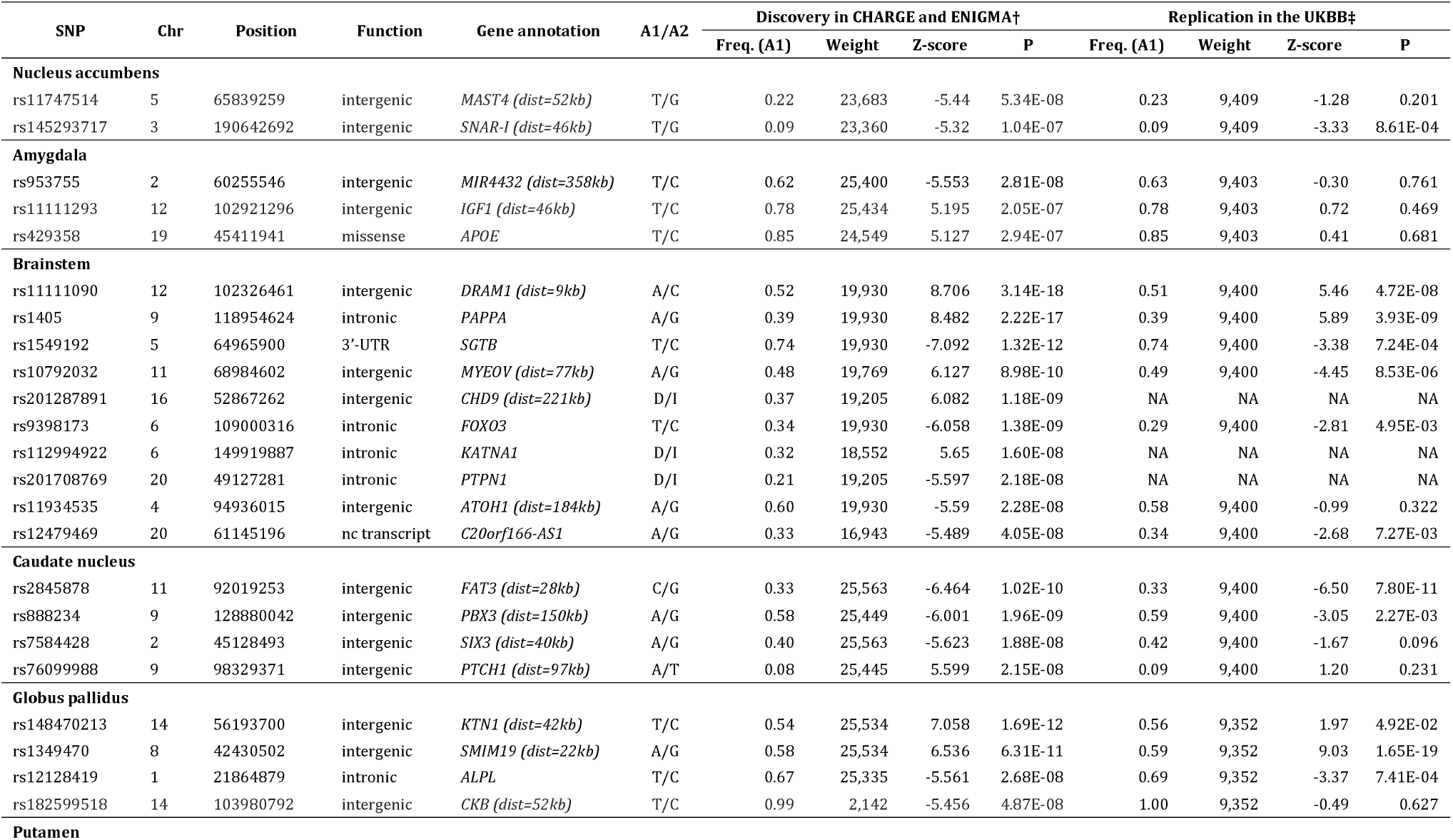
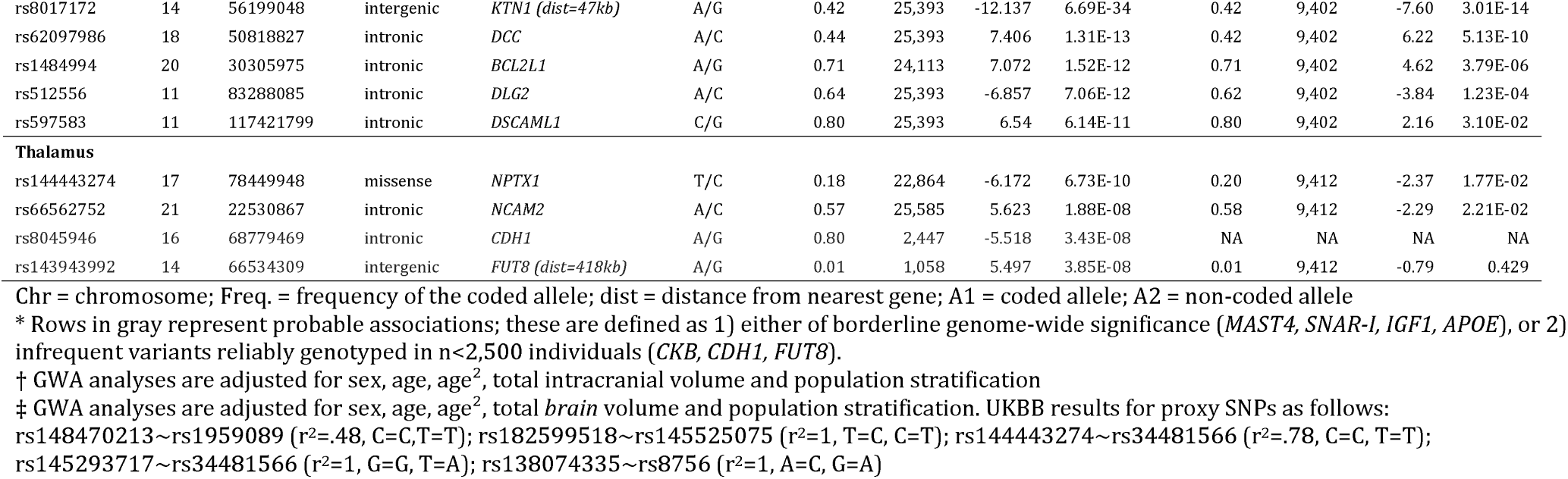
Genome-wide and probable* association results for subcortical brain volumes in the discovery meta-analysis in more than 25,000 Europeans from CHARGE and ENIGMA, and replication results in more than 9,000 Europeans from the UKBB

To functionally annotate our discoveries, we investigated expression quantitative trait loci (eQTL, Table S4) and methylation QTL (meQTL, Table S5) for the 32 candidate loci identified in the discovery analysis, using data from post-mortem brains from the Religious Order Study and the Rush Memory and Aging Project (ROSMAP). We also queried a variety of *cis-* and trans-eQTL datasets in brain and non-brain tissues (further described in the Supplement) for the 32 candidate loci or their proxies (r^2^>0.8), using the European population reference (Table S6). This allowed us to identify 31 additional candidate genes (in addition to the 31 candidate genes within or near the 32 loci carried forward for *in-silico* replication), including one long intergenic non-protein coding RNA, and one microRNA, yielding a final set of 62 candidate genes (Table S7). The details describing the process, whereby specific genes were identified at each locus, can be found in the supplement (see extended results in the Supplementary note).

### Associations with cognitive function and neuropathological phenotypes

We related genetic variation of the 32 variants as well as the expression of our final set of 62 genes influencing subcortical brain volumes to cognitive function and neuro-pathological traits in ROSMAP. We did not find significant associations for individual variants with any investigated trait after Bonferroni correction (P<0.0003), except for the *APOE* variant rs429358, which was, not surprisingly, associated with the presence of neurofibrillary tangles, tau density, β-amyloid load, neuritic plaques, and cognitive decline (Table S8). However, we did find significant associations of dorsolateral prefrontal cortex mRNA expression levels of five candidate genes influencing brainstem, caudate, and putamen volumes (Table S9). These included associations with cognitive function (*KTN1, BCL2L1, SGTB, C20orfl66-ASl, PTCH1*), neuritic plaque presence (*BCL2L1, KTN1*), β-amyloid load (*SGTB, KTN1*), neurofibrillary tangles (*BCL2L1*), and tau density (*BCL2L1*).

### Phenotypic and genetic correlations

We explored both phenotypic and genetic correlations among subcortical volumes, and also the genetic correlations between subcortical volumes and height, MRI-defined hippocampal^17^ and intracranial^18^ volumes, adult height^19^, body mass index^20^, Alzheimer’s disease^21^, general cognitive function^22^, bipolar disorder^23^, and schizophrenia^24^; using linkage disequilibrium (LD) score regression methods^25^ (Figure 2 and Supplementary Table S10). We observed strong phenotypic (P<3.95E-^06^) and genetic (P=0.04-4.5×10^-17^) overlap among all subcortical structures (Figure 2A), consistent with our finding that many of the loci identified have pleiotropic effects on the volumes of several subcortical structures (Table S3).

**Figure 2.**
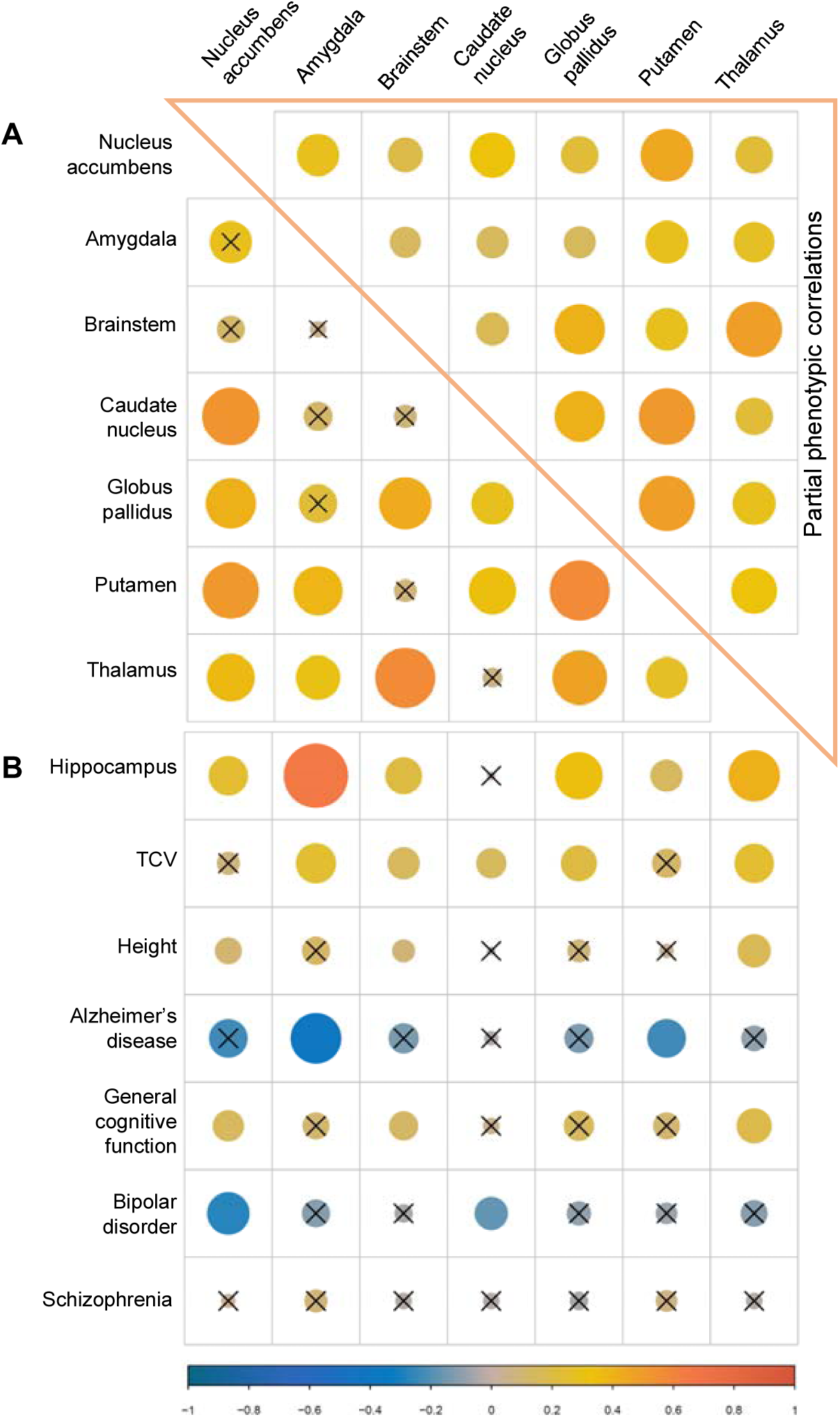
Genetic and phenotypic correlations. In this heat map, the size of the circle is proportional to the strength of correlation (ρ) and the direction is presented in the color label on the bottom; ‘X’ indicates no significant association (p>0.05). **(A)** Partial phenotypic (upper triangle) and genetic (lower) correlations among the subcortical structures included in this report. Partial phenotypic correlations were derived from the subcortical volumes of n=894 participants from the Framingham Heart Study, adjusting for sex, age, age^2^, total intracranial volume and PC1. **(B)** Genetic correlations using LD score regression between subcortical brain volumes and other MRI-derived volumes, anthropometric, and neuropsychiatrie traits.

As expected, we found strong genetic correlations among the nuclei composing the corpus striatum, particularly for nucleus accumbens with putamen (P=1.24×10^-14^), and with caudate nucleus (P=6.92×10^-13^). The genetic architecture of thalamic volume highly overlapped with that of most subcortical volumes, except for the nucleus accumbens. In contrast, there were no significant genetic correlation of the volume of the brainstem with that of most other structures, with the exception of very strong correlations with volumes of the thalamus (P=4.45 ×10^-17^) and the globus pallidus (P=9.20 ×10^-09^).

We also observed strong genetic correlations of smaller amygdala and putamen volumes with increased risk of Alzheimer’s disease, and smaller nucleus accumbens and caudate nucleus volumes with risk of bipolar disorder. Increased general cognitive function was correlated with larger brainstem, thalamic, and nucleus accumbens volumes. Finally, intracranial volume was genetically correlated with larger volumes of subcortical structures, except for the nucleus accumbens and the putamen (Figure 2B).

### Cross-species analysis

To investigate for potential evolutionarily conserved requirements of our gene-set in neurodevelopment, neuronal maintenance, or both, we examined available genetic and phenotypic data from the fruit fly, *Drosophila melanogaster.* Importantly, compared to mammalian models, the fly genome has been more comprehensively interrogated for roles in the nervous system. We found that the majority of candidate genes for human subcortical volumes are strongly conserved in the *Drosophila* genome (66.1%), and many of these genes appear to have conserved nervous system requirements (Table S11). To examine if this degree of conservation was greater than that expected by chance, we leveraged systematic, standardized phenotype data based on FlyBase annotations using controlled vocabulary terms (Table S12). Indeed, 24.1% of the conserved fly homologs are documented to cause “neuroanatomy defective” phenotypes in flies, representing a significant (P=3.9×10’^-3^), nearly two-fold enrichment compared to 12.9% representing all *Drosophila* genes associated with such phenotypes (Table S13).

### Protein-protein interactions

To explore potential functional relationships between proteins encoded by our set of 62 genes, we conducted protein-protein interaction analyses in STRING^26^. Our results revealed enrichment of genes involved in brain-specific pathways (i.e. nervous system development, regulation of neuronal death, neuron projection, axon, neuron part), as well as housekeeping processes (i.e. cell differentiation, apoptosis, kinase binding). Figure 3 shows these protein networks, and the detailed pathways are presented in Table S14.

**Figure 3.**
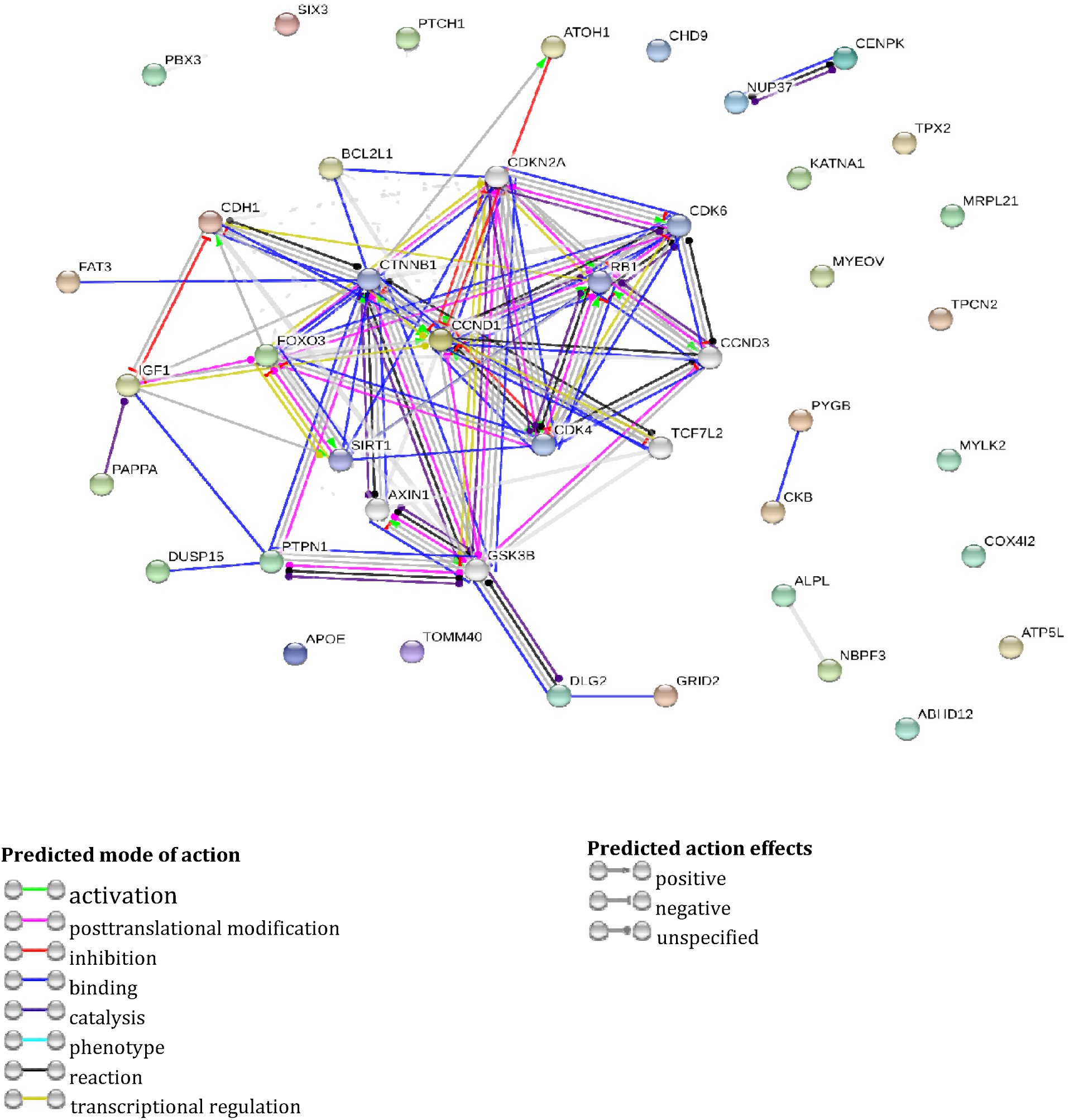
Protein-protein interaction network of 57 genes enriched for common variants influencing the volume of subcortical structures using medium-confidence interaction scores from the human STRING database. The edges represent protein-protein associations, where the edge color indicates the predicted mode of action and the edge shape the predicted action effects (see labels on the bottom]. Colored nodes represent the queried proteins and first shell of interactors (5 maximum], whereas white nodes represent the second shell of interactors (5 maximum).

## DISCUSSION

We undertook the largest GWA meta-analysis of variants associated with MRI-derived volumes of the nucleus accumbens, amygdala, brainstem, caudate nucleus, globus pallidus, putamen, and thalamus; in more than 40,000 individuals from 54 study samples worldwide. Our analyses identified a set of 62 candidate genes influencing the volume of these subcortical brain structures, most of which have well-established roles in the nervous system.

We identified genes implicated in ***neurodevelopmental processes***, including all the candidates influencing the volume of the caudate nucleus. We confirm one locus in llql4.3 near the *FAT3* gene previously associated with the caudate nucleus^13^, where the top variant is an eQTL for the expression of *FAT3* in CD14+ monocytes (Table S6). This gene encodes a conserved cellular adhesion molecule implicated in neuronal morphogenesis and cell migration based on mouse genetic studies^27^. Variants in a locus on 9q33 located 150kb from *PBX3* were also significantly associated with caudate volume. *PBX3* is robustly expressed in the developing caudate nucleus of the non-human primate, *Macaca fuscata*, consistent with a role in striatal neurogenesis^28^. Another locus associated with caudate volume at 2p21 is 40kb proximal to *SIX3*, which encodes a transcriptional regulator with conserved neurodevelopmental roles in both vertebrates and invertebrates^29^. The most significant variant at this locus is associated with CpG sites near active transcription start sites (TSS) harboring *SIX3* in anterior caudate brain tissues (Figure S3.F). Finally, another locus associated with caudate volume was at the 9q22.3 locus, 97kb upstream of *PTCH1*, encoding a receptor for the Sonic Hedgehog (SHH) signaling protein, which was also recently found associated with hippocampal volume^17^. Mutations in *PTCH1* and *SHH* are responsible for a third of medulloblastomas^30^. In addition, dominant mutations in *SIX3, PTCH1*, and *SHH* similarly cause human holoprosencephaly^31^, and their genetic manipulation causes analogous developmental phenotypes in mice^30,32^. Moreover, *SHH* is a direct transcriptional target of *SIX3*^33^, raising the possibility that this pathway also regulates caudate development.

Furthermore, in our GWA of brainstem volume we identified a signal at 4q22,185kb downstream of *AT0H1*, an important gene for neurodevelopment. *AT0H1* encodes an evolutionarily conserved transcriptional regulator of neuronal differentiation, based on studies in numerous animal models^34^. Mice lacking *Mathl*, the *AT0H1* ortholog, show widespread brainstem developmental anomalies^35^, including disruption of medullary and pontine nuclei with roles in respiratory drive^36^. The most significant variant in this locus is also an eQTL for the expression of *SMARCAD1* and *GRID2* in blood cells (Table S6). In mouse experimental models, expression of *Smarcad 1* accompanies neurogenesis^37^; whereas in Lurcher mice, serving as a model for neurodegeneration, mutations in *Grid2* are characterized by brainstem and cerebellar neurodegeneration^38^ resulting in ataxia^39^. We found that variants in *PAPPA* and *IGF1* are associated with the volumes of the brainstem and caudate nucleus, respectively. *PAPPA* encodes a secreted metalloproteinase that cleaves IGFBPs, thereby releasing bound IGF. Although IGF may be beneficial in early- and midlife (i.e. higher levels are associated with larger brain volumes and a lower risk of Alzheimer’s disease^40^); its effects may be detrimental during aging, and studies of PAPPA similarly support antagonistic pleiotropy. Low circulating PAPPA levels are a marker for adverse outcomes in human embryonic development^41^, but in later life, higher levels have been associated with acute coronary syndromes and heart failure^42,43^. Similarly, *Pappa* knockout mice show dwarfism but reduced age-related degeneration and increased longevity^44^.

In screening for variants associated with globus pallidus volume, we identified additional genes involved in neurodevelopment. One was an intronic variant in *ALPL*, associated with CpG sites near enhancers in the gene and transcription sites in *NBPF3* (Table S5 and Figure S3.1). *ALPL* encodes an alkaline phosphatase that mediates bone mineralization, regulates cell migration, neuronal differentiation early during development, and post-natal synaptogenesis in transgenic mouse models^45^. Recent reports suggest that ALPL helps propagate the neurotoxicity induced by tau^46^, and its activity increases in Alzheimer’s disease^47^ and cognitive impairment^48^. *NBPF3* belongs to the neuroblastoma breakpoint family, which encodes domains of the autism- and schizophrenia-related DUF1220 protein^49^.

Genes influencing the volume of the thalamus, a relay hub for electrical impulses travelling between subcortical structures and the cerebral cortex, were related to ***synaptic signaling pathways.*** We found a missense variant in *NPTX1*, a gene expressed in the nervous system which restricts synapse plasticity^50^, and induces β-amyloid neurodegeneration in human and mouse brain tissues^51^. We also identified an intronic variant in *NCAM2*, encoding a protein involved in olfactory system development^52^, levels of which are lower in hippocampal synapses of Alzheimer’s disease brains^53^, possibly contributing to synapse loss in Alzheimer’s disease.

Additionally, the identified variant at the 3’-UTR of *SGTB* for the brainstem was a robust eQTL for the expression of SGTB in cerebellum, visual cortex (Table S6), and dorsolateral prefrontal cortex (Table S4). Experimental rat models showed that βSGT, highly expressed in brain, forms a complex with the cysteine string protein and heat-shock protein cognate (CSP/Hsc70) complex to function as a chaperone guiding the refolding of misfolded proteins near synaptic vesicles^54^. Other experimental studies in the nematode worm, *C. elegans*, showed that the genetic manipulation of the ortholog, *sgt-1*, suppresses toxicity associated with expression of the human β-amyloid peptide^55^. Other genes involved in synaptic signaling are *CHPT1* (brainstem), involved in phosphatidylcholine metabolism in the brain, and *DLG2* (putamen), encoding an evolutionarily conserved scaffolding protein involved in glutamatergic-mediated synaptic signaling and cell polarity^56^ that has been associated with schizophrenia^57^, cognitive impairment^58^, and Parkinson’s disease^59^.

Other identified variants point to genes involved in *autophagy and apoptotic processes*, such as *DRAM1* and *F0X03*, both related to brainstem volumes. *DRAM1* encodes a lysosomal membrane protein involved in activating TP53-mediated autophagy and apoptosis,^60^ and mouse models mimicking cerebral ischemia and reperfusion have found that inhibiting the expression of *DRAM1* worsens cell injury^61^. The most significant variant located 9Kb downstream from *DRAM1* was also associated with a CpG site proximate to active TSS upstream of that gene in several mature brain tissues (Table S5 and Figure S3.B). *F0X03* has been recently identified as pivotal in an astrocyte network conserved across humans and mice involved in stress, sleep, and Huntington’s disease^62^, and has been related to longevity^63^. In *Drosophila*, a *F0X03* ortholog regulates dendrite number and length in the peripheral nervous system^64^, and in the zebrafish, *Danio rario, Foxo3a* knockdown led to apoptosis and mispatterning of the embryonic CNS^65^.

Finally, some of the genes we identified have been implicated in *axonal transport.* Our results confirm an association between variants in the 13q22 locus with putamen and globus pallidus volumes as previously reported^13,66^. The most significant variant (rs8017172) is a robust eQTL for *KTN1* in peripheral blood cells (Table S6). This gene encodes a kinesin-binding protein involved in the transport of cellular components along microtubules^67^, and impairment of these molecular motors has been increasingly recognized in neurological diseases with a subcortical component^68^. The 5q12 locus, associated with nucleus accumbens volume in the combined analysis, lies 53kb upstream from *MAST4*, which encodes a member of the microtubule-associated serine/threonine kinases. This gene has been associated with hippocampal volumes^17^ and juvenile myoclonic epilepsy^69^,and it appears to be differentially expressed in the prefrontal cortex of atypical cases of frontotemporal lobar degeneration^70^. In *Drosophila*, the knockdown of a conserved *MAST4* homolog enhanced the neurotoxicity of human tau^71^, which aggregates to form neurofibrillary tangle pathology in Alzheimer’s disease.

Overall, the loci identified by our study pinpoint candidate genes not only associated with human subcortical brain volumes, but also reported to disrupt invertebrate neuroanatomy when manipulated in *Drosophila* and many other animal models. This is consistent with the results observed in protein-protein networks. Thus, our results are in line with the knowledge that the genomic architecture of central nervous system development has been strongly conserved during evolution. Further elaboration of the biological pathways associated with the genes not discussed in the main text may be found in the Supplementary note (see extended results).

Our findings derived from genetic correlations support earlier observations that amygdala volume is reduced in Alzheimer’s disease patients^72^ and in carriers of the Alzheimer risk enhancing £4 variant of the APOE gene^73^. Interestingly, one of the top signals related to the amygdala was one of the two variants that determines the *APOE ε4* isoform (rs429358). In line with our findings, other studies have described smaller putamen volumes in Alzheimer’s disease^74^, or smaller accumbens and caudate nuclei in patients with bipolar disorder^75,76^. Notably, higher general cognitive function was correlated with larger brainstem, thalamus, and nucleus accumbens, highlighting the integrative role of these brain structures in cognition.

In conclusion, we describe multiple genes associated with the volumes of MRI-derived subcortical structures in a large sample, leveraging diverse bioinformatic resources to validation and follow-up our findings. Our analyses indicate that the variability of evolutionarily old subcortical volumes of humans is moderately to strongly heritable, and that their genetic variation is also strongly conserved across different species. The majority of the variants identified in this analysis point to genes involved in neurodevelopment, regulation of neuronal apoptotic processes, synaptic signaling, brain homeostasis, and susceptibility to neurological disorders. We show that the genetic architecture of subcortical volumes overlaps with that of anthropometric measures and neuropsychiatric disorders. We have focused on the discovery of common and less frequent variants, but further efforts to also reveal rare variants and epigenetic signatures associated with subcortical structures will provide an even more refined understanding of the underlying mechanisms involved. In summary, our findings greatly expand current understanding of the genetic variation related to subcortical structures, which can help identify novel biological pathways of relevance to human brain development and disease.

